# HPV16 neutralizing monoclonal antibodies show evidence for common developmental pathways and public epitopes

**DOI:** 10.1101/2025.03.31.646278

**Authors:** Joseph J. Carter, Nicholas K. Hurlburt, Erin M. Scherer, Suruchi Singh, Justas V. Rodarte, Robin A. Smith, Peter Lewis, Rachel Kinzelman, Jacqueline Kieltyka, Madelyn E. Cabãn, Gregory C. Wipf, Marie Pancera, Denise A. Galloway

## Abstract

Antibodies to human papillomavirus (HPV) primarily recognize surface exposed residues on five loops of the major capsid protein (L1) that vary significantly among HPV types. We determined which loops were required for neutralization for 70 HPV16 specific human monoclonal antibodies (mAbs) cloned from participants who received an HPV vaccine, and we describe molecular features of those antibodies.

Chimeric HPV16 pseudovirus (cpsV), each having one surface loop bearing multiple amino acid substitutions, were used to determine neutralization specificity. The HPV16-FG-loop was the loop most frequently required for neutralization (44 of 70, 62.9%), however, all other surface loops were used for neutralization by multiple mAbs: HI (13, 18.6%), DE (15, 21.4%), EF (six, 8.6%), BC (four, 5.7%). Antibodies that required multiple loops were common (17, 24.3%). Three mAbs (4.3%) required sequences on the c-terminus of L1 and for another three mAbs the neutralization specificity could not be determined.

Two types of mAbs appeared to be overrepresented: ten mAbs used V_H_ 2-70 IGHV paired with V_L_ λ1-40, having characteristic mutations in complementarity determining region two (CDRL2). Cryogenic electron microscopy (Cryo-EM) revealed that two of these antibodies bound five Fabs per pentamer interacting with all five L1-surface loops. The other type of mAbs that appeared to be overrepresented were ten mAbs using V_H_ 4-34, seven of which also used D_H_3-16*02 with conserved CDRH3 sequences. Cryo-EM for one of these mAbs, that required the FG-loop for neutralization, was shown to bind one Fab per pentamer at the apex, interacting with the DE- and FG-loops, with sequences of the Fab CDRH3 inserted between the DE- and FG-loops from two protomers. These two types of mAbs were found repeatedly in the four participants suggesting that these antibodies shared developmental pathways and bound to similar immunodominant epitopes on the virus.

**Highlights:** Most human mAbs recognized L1 surface loops but three of 70 recognized sequences on the C-terminal arm of L1

Some antibodies induced by HPV vaccination follow shared developmental pathways.

Human monoclonal antibodies using V_H_ 2-70/V_L_ λ1-40 were found in all participants and bound with at a stoichiometry of five Fabs per capsomer.

Human monoclonal antibodies using the diversity gene segment D3-16*02 were found in all participants and one Fab was shown to bind with a stoichiometry of one Fab per capsomer.

**In brief:** A panel of 70 HPV16 specific human monoclonal antibodies (mAbs), cloned from memory B cells or plasmablasts following HPV vaccination, was characterized by determining the surface loops of the major capsid protein (L1) required for neutralization and examined for shared molecular features. All five L1 loops were found to be used for neutralization by one or more antibodies, but the most frequent target of these antibodies was the FG loop followed by the HI and DE loops. Ten antibodies paired the heavy chain variable gene V_H_ 2-70 with the light chain variable gene V_L_ λ1-40 and these antibodies had conserved mutations in the CDRL2 region of V_L_ λ1-40. Mutating the CDRL2 back to the predicted germline sequence significantly reduced neutralization. Cryo-EM analysis of two V_H_2-70/V_L_λ1-40 mAbs showed five Fabs binding per L1 pentamer and a conserved epitope with Fabs interacting with all five variable loops across two adjacent protomers. Seven other mAbs had a heavy chain composed of the variable region V_H_4-34 with the diversity gene D3-16*02 resulting in the sequence motif WSGYR in the CDRH3. Mutation of that sequence to alanine ablated HPV16 neutralization activity. A cryo-EM structure of one of these antibodies showed one Fab binding the pentamer apex with the WSGYR motif inserting between three loops from two protomers. Antibodies with paired V_H_ 2-70/V_L_ λ1-40 and the antibodies with CDRH3 containing the WSGYR sequence, were found in all four study participants suggesting that such antibodies may be commonly elicited following HPV vaccination.

## Introduction

Sexually transmitted human papillomavirus (HPV) infections are common and generally benign but occasionally result in malignant transformation of infected cells. HPVs classified as “high risk” infect mucosal epithelium and cause virtually all cervical cancers ^1^, a significant proportion of other anogenital cancers and head and neck cancers ^2,3^. HPV16 is the HPV type most frequently found in cancers ^4^, accounting for approximately 50% of cervical cancers and greater than 90% of oropharyngeal cancers ^3^. Current vaccines, when administered prior to the initiation of sexual activity, provide sustained protection from infection and disease ^5,6^. Currently, two doses of vaccine are recommended ^7^ for children under 15 years old, and there is growing evidence that even a single dose of an HPV vaccine is protective ^8,9^. We were interested in better characterizing the B cell response elicited by such an effective vaccine.

The HPV virion is composed of an approximately eight kb circular double stranded DNA genome, encapsidated by major (L1) and minor (L2) capsid proteins that assemble into a T=7 icosahedral particle composed of 72 pentameric capsomers ^10^. Twelve pentamers are at five-fold axes with 60 occurring at six-fold or hexavalent axes. The c-terminus of each L1 connects capsomers by an extension, referred to as an arm, that bridges the intercapsomer canyon, forming disulfide bonds with the neighboring pentamer and returning to the parent capsomer ^11^. Although the C-terminal arm has been shown to be immunogenic, ^12–14^ most antibodies target sequences encoded by five surface exposed loops (BC, DE, EF, FG and HI) ^10,15–17^ and because the amino acid sequences that form these loops vary by type, antibody responses tend to be type-specific or weakly cross-reactive ^18,19^.

With most anti-HPV antibodies recognizing the native conformation of L1, a combination of loop-swaps, site specific mutagenesis and cryogenic electron microscopy (cryo-EM) have been used to discover how antibodies bind virus and the residues on capsids important for recognition and neutralization ^15–17,20–24^. Two types of antibodies have been repeatedly found ^21,23,25^. The majority of antibodies described interact with residues on multiple loops, often including the FG loop. These antibodies which include the well-studied murine mAb H16.V5, bind 5:1 (5 Fabs per pentamer) in a manner described as top fringe binding ^21–23^. The other type of antibody described multiple times, binds 1:1 (1 Fab per pentamer) at the apex, contacting residues on the DE and FG loops ^21,23,25,26^. Antibodies recognizing sequences on a C-terminal arm of L1 bind in the canyon around hexavalent capsomers ^12,13^ but only one such antibody has been described.

Characterization of antibody sequences induced by vaccination or following viral infection has led to a greater understanding of virus – host interactions, insights into the evolution of protective antibody responses and suggested new approaches for vaccine design ^27–33^. Following vaccination or infection, antibodies from different people often share similar characteristics suggesting that, from the immense array of naïve B cells, a limited number of germline sequences are selected that follow similar pathways of affinity maturation and bind to shared, or public, epitopes. These recurrent antibodies may be neutralizing and, occasionally, broadly neutralizing ^34^.

Few studies have examined the molecular features of antibodies important for recognition and neutralization of HPVs ^13,22,35,36^. Our studies on vaccine induced B-cell responses resulted in the molecular cloning of a large number of human mAbs from HPV16 specific memory B-cells and plasmablasts ^37^. In this study we map L1 loops required for neutralization by 70 human mAbs, identify two types of mAbs that were found in all four study participants and describe the structural basis for their strain specific neutralization by cryo-EM. Our findings suggest that individuals develop antibodies with common features that bind similar, perhaps public, epitopes.

## Results

### Surface loops of HPV16 required for neutralization

The mAbs studied here (n=70), were cloned from memory B cells or plasmablasts, isolated from four study participants who had received the quadrivalent Gardasil® vaccine (4vHPV) at the recommended schedule of 0, two and six months with an additional off-label vaccine booster dose at month 24 ^38^.

Human mAbs (S1 Table cross-references previous mAb nomenclature) were initially tested for HPV16 binding and neutralizing activity. With few exceptions, it was found that binding antibodies were neutralizing (S1 Fig) with most having IC_50_s between one and ten pM (S2 Fig). No mAb tested cross-reacted with HPV18 in binding assays (not shown). To identify residues required for neutralization, cpsV were created in which each of the five surface loops or a region of the c-terminal of L1 of HPV16 were substituted with sequences from HPV35, HPV18 or HPV2 (Fig 1A). Due to the length of the FG loop, cpsVs were created for amino-terminal and carboxyl-terminal halves of this loop (FGa and FGb). We were not able to study all variable surface residues by this method because substitution of some residues (such as the C-terminal portion of the HI loop) did not result in infectious psV (not shown). All cpsV reported in this study were infectious allowing us to directly assess sequences required for neutralization. HPV31 psV (psV31) was included in this screen to determine if mAbs were cross-reactive as HPV31 was not included in the 4vHPV vaccine.

**Fig 1.**
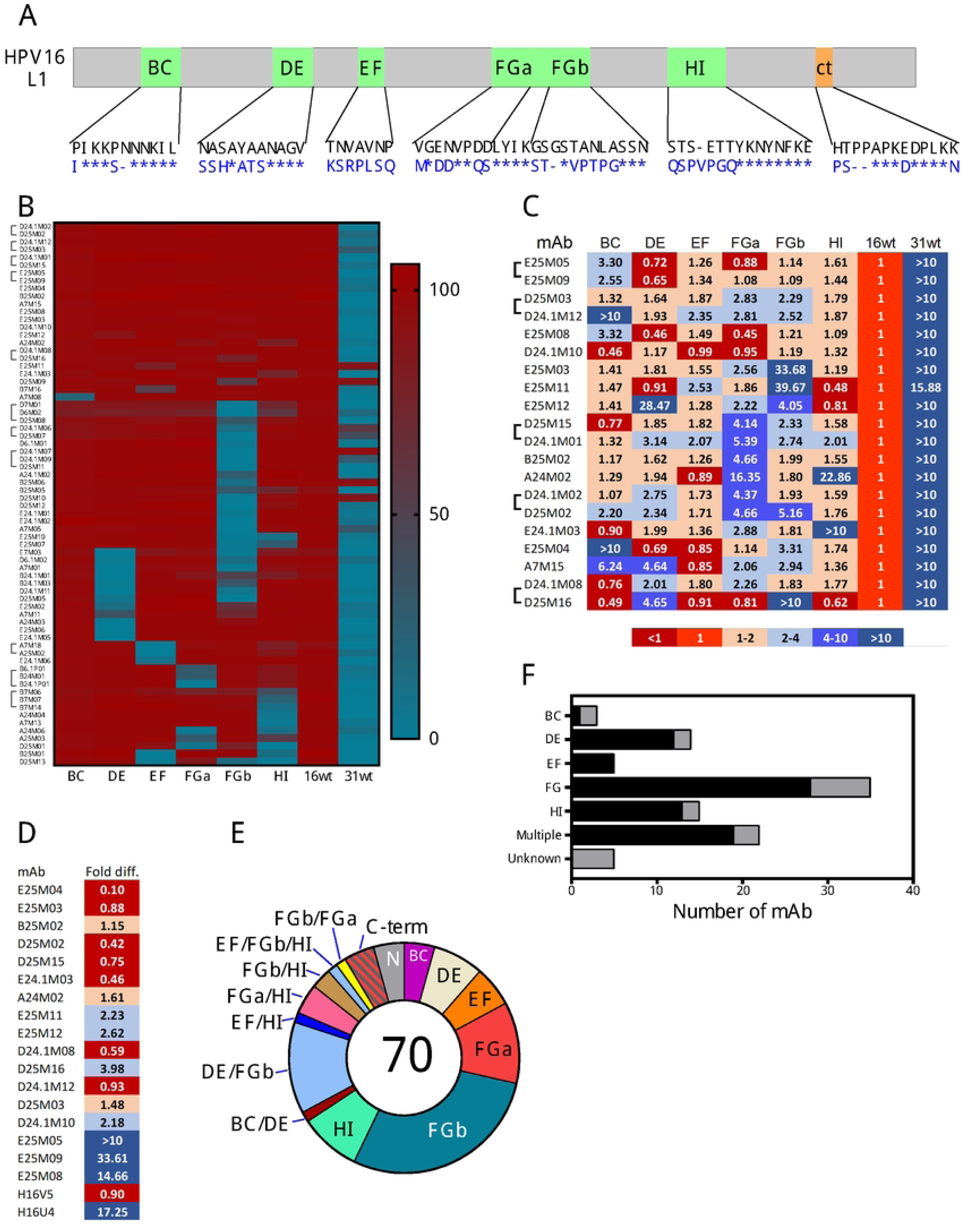
Delineation of sequences required for neutralization by human mAbs. A) Diagram of L1 protein with surface loops (green) and c-terminal variable region (orange) indicated. Loop and c-terminal sequences are shown with amino acid substitutions (blue letters), inserts and deletions. Asterisks indicate no change from HPV16 sequence (black letters). B) The percent neutralization of 70 mAbs (on left) against psVs (listed on bottom) are represented in a heat map after manual clustering. Brackets indicate antibody clonotypes. C) Select antibodies were tested in a secondary neutralization screen against psVs listed on top. IC_50_ were calculated and the log fold change in IC_50_ of cpsV compared with psV16 are represented by color. D) Similar to panel (C) however only cpsV with mutations on the c-terminus were tested along with psV16. Colors represent fold changes as in C. E) The specificity for each of 70 mAbs, based on neutralization experiments represented above is summarized in a pie chart. F) Graph depicting the number of mAbs that required each surface loop (bottom), multiple loops or the c-terminus detected in either the primary (black bar) or secondary (gray bar) neutralization screens.

To determine which sequences of HPV16 L1 were required for neutralization by mAbs, assays were conducted with cpsV having substitutions on one of five surface loops (Fig 1A). To conserve reagents, a concentration of antibody 100-fold higher than the IC_50_ for HPV16 (psV16) with no amino acid substitutions was used for the initial screen (Fig 1B, S2 Table). Twenty mAbs neutralized all cpsV by greater than 55%. These mAbs were serially diluted for a secondary screen to determine IC_50_s against each cpsV (Fig 1C). Six mAbs neutralized all cpsV with IC_50_s less than four-fold greater than for psV16. These antibodies were subsequently tested in neutralizing assays against cpsV that had amino acid substitutions on the C-terminus (psV16-c-term) (Fig 1D). Neutralizing epitopes for each mAb were considered to include a loop if a mAb failed to neutralize a particular cpsV by more than 55% in the first screen, or had an IC_50_ at least four-fold higher than psV16 in secondary screens. Most mAbs (54 of 70, 77.1.%) were type specific and failed to neutralize closely related psV31 more than 10%, five mAbs (7.1%) effectively neutralized (>88%) psV31 and 11 (15.7%) had intermediate levels (between 10% and 60%) (Fig 1B). There were 15 different patterns of neutralization (Fig 1E). Four mAbs required each loop for neutralization, 17 (24.3%) mAbs required two or more loops for neutralization and three (4.3 %) mAbs were able to neutralize all cpsV tested (epitope undefined). The majority of neutralizing mAbs (44, 62.8%) required loop FG (FGa and/or FGb) as part of the neutralizing epitope (Fig 1F). The next most frequently required loops for neutralization were DE (15, 21.4%) and HI (13, 18.6%). The EF loop was required for neutralization by six mAbs (8.6%), the BC loop by four mAbs (5.7%) and three antibodies (4.3%) required the c-terminus. Thus, most HPV16 binding mAbs were potent neutralizing antibodies that bound residues on the FG loop for neutralization, however, there were many different specificities, and all loops were required by some mAbs. Clonotypes (defined below, bracketed in Fig 1 B, C, D) had the same pattern of neutralization in most instances.

### Antibodies with HPV16-HPV31 cross-neutralization activity

HPV31 is closely related to HPV16 and people vaccinated for HPV16 have partial protection from infection with HPV31 ^39^. We included psV31 in neutralization assays to discover mAbs that might account for cross-protection. Six mAbs were defined here as cross-reactive as they could prevent in vitro infection with both psV16 and psV31 (at least 55%). Five of these antibodies required the HPV16 FGb loop for neutralization activity in the initial screen (Fig2 A), E25M11 was found to require FGb in the second screen (Fig1 C). A previous report had found HPV31-HPV35 cross-reactive epitopes on loops DE and FG ^16^, however these 6 HPV31 cross-reactive antibodies did not require the DE loop for neutralization in our assays. One cross-reactive antibody (D24.1M07) was a member of a clonotype consisting of four antibodies that had variable levels of HPV31 cross-reactivity (Fig 2B, C). To determine which sequences accounted for the increased cross-reactivity, heavy-light chain-swap mAbs were generated using the cross-reactive mAb D24.1M07 and non-cross reactive mAb D25M11. The swapped mAbs neutralized psV16 equally well (Fig 2D) however, the swapped mAb having a D24.1M07 light chain neutralized psV31 approximately 10 fold more effectively (Fig 2D). In summary, all cross-reactivity mAbs required the FGb loop for neutralization and, for one mAb, cross-neutralization of psv31 was primarily due to the light chain.

**Fig 2.**
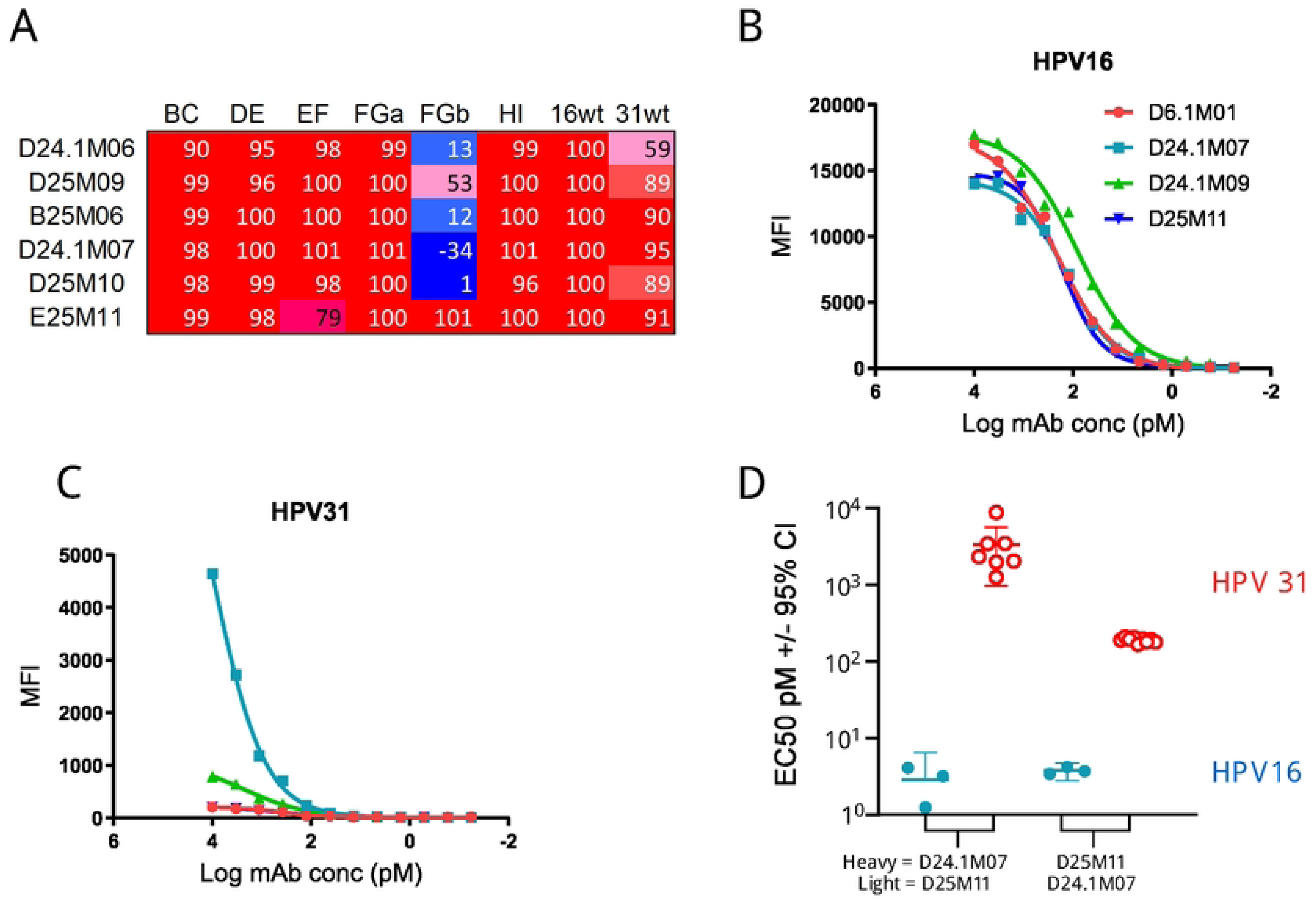
HPV31 cross-reactive mAbs. A) The percent neutralization determined in the primary screen against each psV are shown for six mAbs with the highest psV31 cross-reactivity. Four Mabs from the same clonotype bound HPV16 similarly (B) but had variable binding to HPV31 (C). Chimeric mAbs were created by swapping the heavy and light chains for antibodies D24.1M07 and D25M11 and used in neutralization assays (D). Each data point represents a neutralization EC50 (vertical axis) calculated from a titration series. MAbs having swapped heavy and light chains, indicated on bottom, tested against psV16 (red filled circles) and psV31 (blue empty circles) are shown.

### Antibody features associated with neutralization specificity

To avoid biasing our analysis of antibody gene usage we collapsed our set of antibodies to include only 54 unique clonotypes derived from memory B cells. Clonotypes were defined here as antibodies from the same individual, with the same V_H_ and V_L_, the same length CDR3 regions and greater than 70% amino acid sequence identity in the CDRH3.

To determine if these mAbs shared genetic features that corresponded to recognition of specific L1 loops, we focused on antibodies that employed the IGHV2-70 or IGHV3-34 heavy chain variable gene segments (V_H_) as they were the most frequently used V_H_ in the collapsed set (8/54, 14.8%, for both, Fig 3A). Another antibody gene sequence that appeared to be over-represented was IGkV1-39, with 15 clonotypes (19 mAbs, Fig 3A). However, these antibodies did not share a common neutralization pattern, paired with eleven different IGHVs and had few somatic mutations in common (not shown).

**Fig 3.**
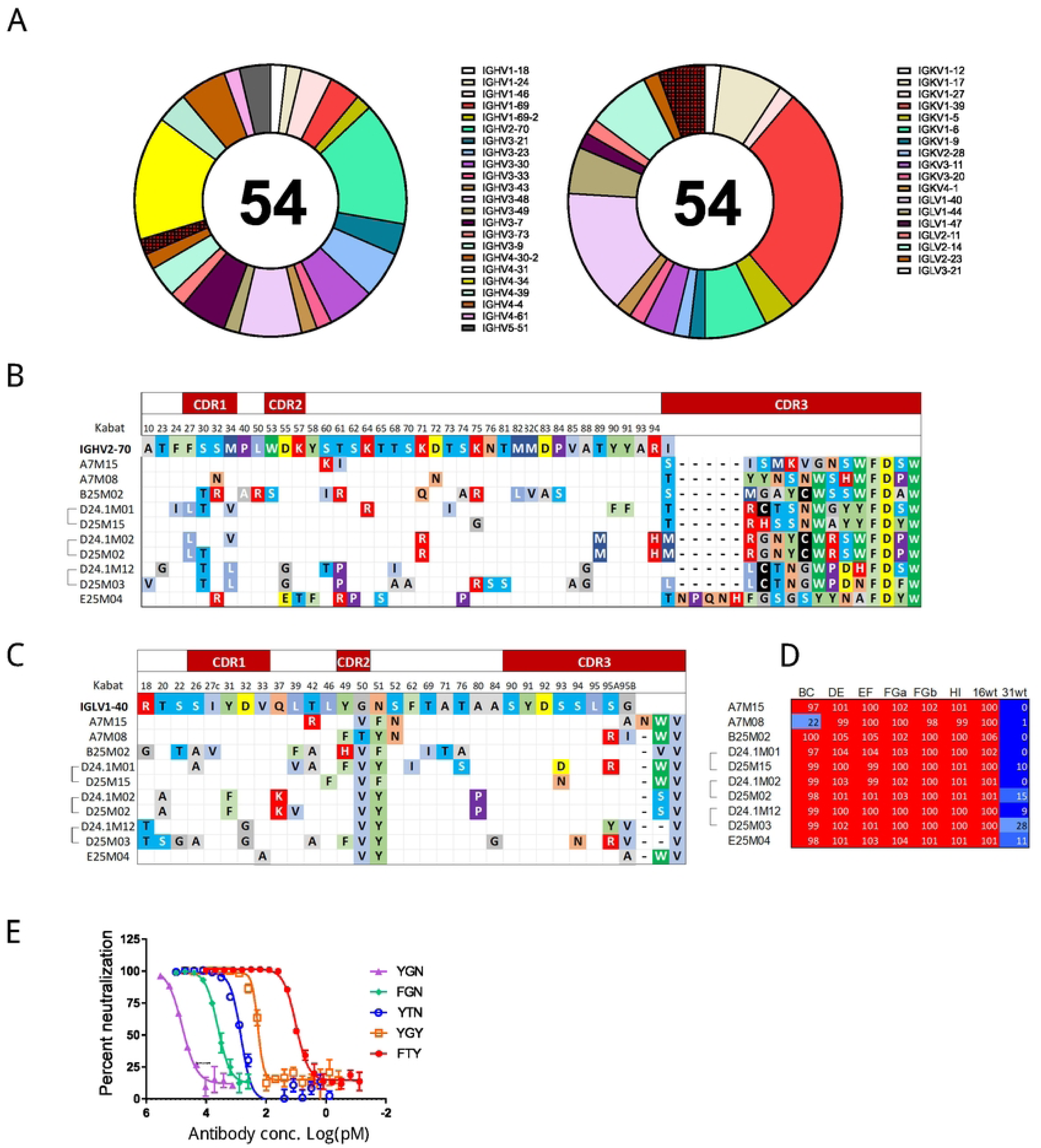
Comparison of mAbs using VH2-70 and VL1-40. A) The number of each IGHV (left) and IGLV (right) gene segment used by the mAbs are depicted in donut graphs with the names of each variable gene segment listed. B) Alignment of mAb sequences using V_H_2-70 that paired with Vλ1-40with the germline V_H_2-70 gene sequence with kabat numbering. Only residues that differed from the reference sequence are shown. All residues in the CDRH3 region are shown. Dashes indicate a gap and empty cells indicate a residue the same as the germline. C) Alignment of the light chain sequences with germline Vλ1-40 that were paired with V_H_2-70. Only residues that differed from the reference are shown. Dashes and empty cell are the same as above. D) The percent neutralization for 10 antibodies using V_H_2-70 paired with V_L_λ1-40 against psVs. E) Substitution mutations reverting putative somatic mutations to predicted germline were made to the CDRL2 region of A7M08, the light chains were co-expressed with the mature heavy chain and resulting antibodies tested for psV16 neutralization. The graph is a representative psV16 neutralization experiment using antibodies with CDRL2 substitution mutations indicated (triplicate tests). Error bars are the standard deviation of triplicate titrations.

Among the eight mAbs that used V_H_2-70, seven were paired with lambda light chain variable region (V_L_) λ1-40 whereas this V_L_ paired with only one other V_H_ among 46 clonotype (Fisher’s exact p < 0.0001). These antibodies tended to have similar length CDRH3 regions (Fig 3B). Despite these similarities, the CDRH3 region sequences were quite diverse (Fig 3B). Evidence for convergent evolution was seen in the V_L_λ1-40 CDRL2 regions (Fig 3C) where all seven unique clonotypes (10 mAbs) had characteristic CDRL2 mutations, resulting in substitution of amino acids G with V at position 51 (all but one) and N52F or N52Y. The mAb that paired V_L_λ1-40 with a different V_H_ did not have similar changes (not shown). We did not identify evidence for convergent sequence changes elsewhere in these antibodies (Fig 3 B,C).

Four V_H_2-70/V_L_λ1-40 mAbs required the BC loop either in the primary (mAb A7M08) or secondary screen (mAbs A7M15, E25M04, D24.1M12 (S2,3 Tables)). The other six antibodies neutralized all cpsV in the primary screen but did not have a consistent pattern of neutralization in secondary screening (S2,3 Tables). Thus, while there were some similarities between these antibodies in neutralization assays, they did not have the same neutralization profile as might be expected based on the lack of sequence homology in the CDR3 regions (Fig 3B).

To evaluate the importance of amino acid substitutions in the CDRL2, the FTY sequence of the mature A7M08 antibody was substituted with the sequence of the predicted germline V_L_λ1-40 gene segment (YGN). The resulting antibody, created by coexpression of the mature A7M08 heavy chain with the light chain having the predicted germline CDRL2, showed decreased neutralization potency with an increased IC_50_ for psV16 by more than three logs compared with mature A7M08 (Fig 3E, S3 Table). To measure the relative importance of each amino acid, germline residues were substituted with the mature sequence found in A7M08. Both the Y49F and G50T substitutions substantially improved neutralization activity compared with the germline sequence (IC_50_s = 4270 +/- 1028 and 908 +/- 230 pM respectively). The N51Y mutation showed the greatest improvement in neutralization compared to the predicted germline sequence with an IC_50_ of 198 +/- 78 pM (Fig 3E). The germline CDRL2 was substituted into six other V_L_λ1- 40 antibodies, and coexpressed with their mature cognate heavy chains (Table S3). Four of the antibodies (A7M15, D24.1M01, D24.1M02, E25M04) shared the pattern seen for A7M08 where substitution to the predicted germline (YGN) sequence significantly increased IC_50_s relative to the mature antibody. Although the pattern was the same for these four antibodies the magnitude of the change was not as great as for A7M08 ranging from 5.3-fold for D24.1M01 to 39.7-fold for E25M04.

Unexpectedly, the IC_50_ for two antibodies (B25M02 and D25M03) were not significantly changed when the CDRL2 was changed to the presumed germline sequence. Similar results were obtained when these antibodies were tested in HPV16L1 binding assays (not shown). To investigate the importance of the CDRL2 for B25M02 neutralization we created an antibody with the mature heavy chain paired with predicted germline V_L_λ1-40. Surprisingly, this antibody had potent neutralizing activity (IC_50_ 30.7 pM +/- 8.2) however, substituting in just the mature CDRL2 (_49_HVF_51_) significantly improved neutralization (IC_50_ 11.2 pM +/- 3.0) (Fig S3). These data confirmed the importance of CDRL2 for the neutralization activity of V_H_2-70 /V_L_λ1-40 antibodies.

### V_H_2-70/V_L_λ1-40 mAbs bind to all variable loops

To determine if V_H_2-70/V_L_λ1-40 mAbs share common binding epitopes and to resolve any key binding features of this class of antibodies, we used cryo-EM to determine the structure of the antigen-binding fragment (Fab) to the HPV16 L1 pentamer. We first chose to look at the binding of D24.1M01, because its neutralization pattern was typical for V_H_2-70/V_L_λ1-40 mAbs, as it failed to neutralize any cpsV in the first screen (Fig 3D). We obtained a cryo-EM reconstruction to 3.0 Å (Fig 4A, Table S4, Fig S4). The structure revealed five Fabs bound to the L1 pentamer at the interface between two protomers (Fig 4B). Each Fab interacts with all five variable loops, the DE- and EF-loops of protomer 1 (P1) and the BC-, FG-, and HI-loops of protomer 2 (P2) (Fig 4C). The Fab binds the two protomers with a total buried surface area (BSA) of 1,745 Å^2^, with 840 Å^2^ from the heavy chain and 905 Å^2^ from the light chain (Fig S5). All six CDRs are involved in binding as well as the framework 3 (FR3) of the light chain. On each L1 protomer, the total BSA is 1,838 Å^2^, with 434 Å^2^, 81 Å^2^, 346 Å^2^, 592 Å^2^, and 385 Å^2^ from the BC-, DE-, EF-, FG-, and HI-loops, respectively (Fig S6).

**Fig 4.**
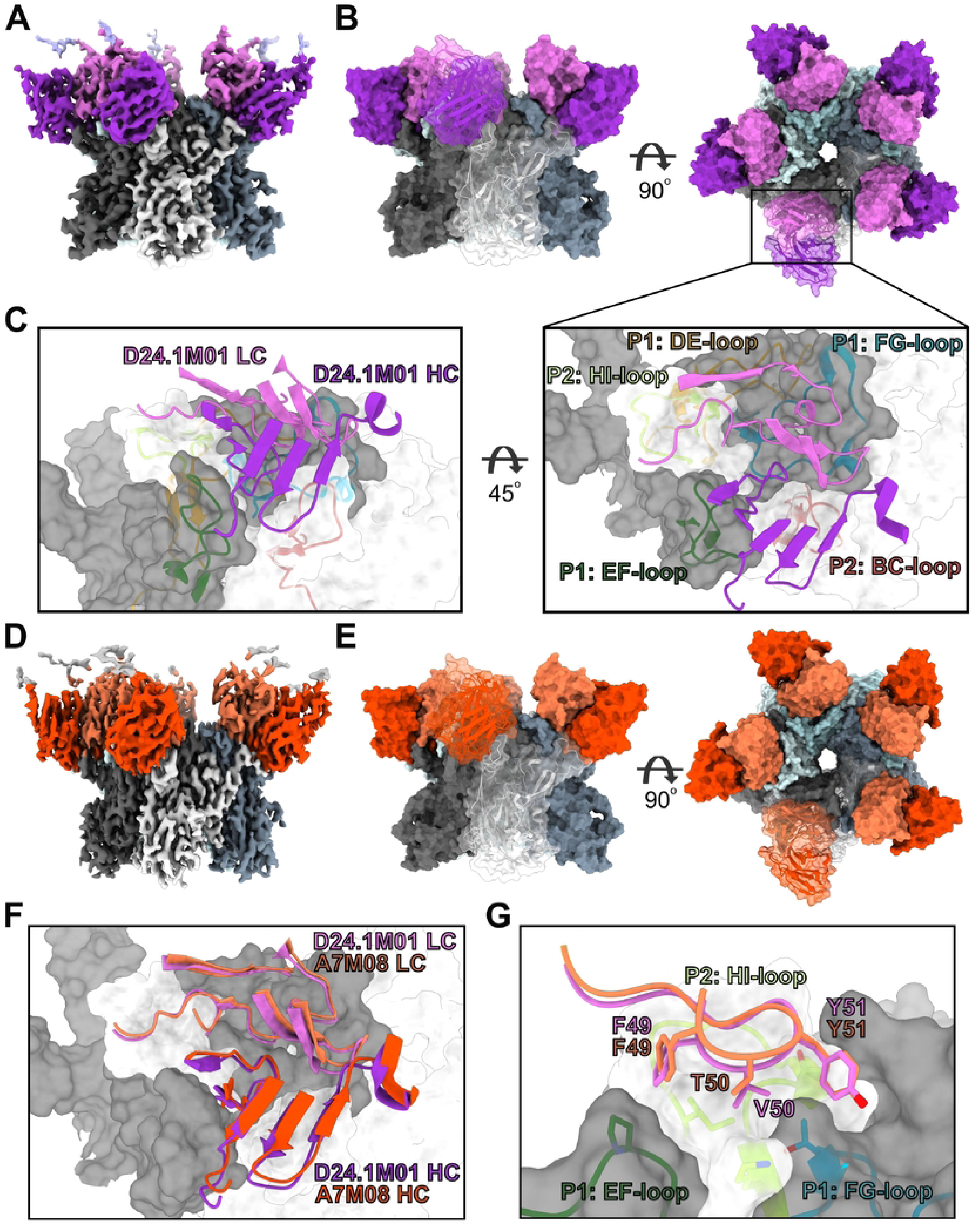
Structures of VH2-70/Vλ1-40 mAbs D24.1M01 and A7M08. A) Cryo-EM map of D24.1M01 Fab bound to the HPV16 L1 pentamer. L1 protomers are colored in shades black, white, and blue. The heavy and light chains of D24.1M01 are colored dark purple and light purple, respectively. B) A surface representation of the structure in two orientations. One protomer and one Fab is shown with a transparent surface with a cartoon representation. C) Cartoon representation of the D24.1M01 CDRs and the footprint on L1. Each L1 variable loop is shown and assigned to a protomer, P1 or P2. D) Cryo-EM map of A7M08 Fab bound to the HPV16 L1 pentamer. L1 protomers are colored in shades black, white, and blue. The heavy and light chains of D24.1M01 are colored dark orange and light orange, respectively. E) A surface representation of the structure in two orientations. One protomer and one Fab is shown with a transparent surface with a cartoon representation. F) Alignment of CDRs from D24.1M01 and A7M08. G) Structure of matured CDRL2 of both D24.1M01 and A7M08 showing the aliphatic surface on L1 that they bind to.

To better understand the role of maturation of the CDRL2 and the difference in binding of these V_H_2-70 /V_L_λ1-40 mAbs, we also chose to determine the structure of A7M08 as it was the one mAb of this group that failed to neutralize cpsV with changes in the BC region (Fig 3D). A7M08 Fab bound to the HPV16 L1 using cryo-EM to a resolution of 2.69 Å (Fig 4D,E, Table S4, Fig S4). The structure showed a nearly identical binding epitope as D24.1M01 (Fig 4F). An alignment of the two structures, including the L1 pentamer and each Fab, has a root mean square deviation (RMSD) of 0.774 Å over 2050 Cα atoms.

A7M08 Fab aligned to D24.1M01 Fab has an RMSD of 0.510 Å over 204 Cα atoms. The total BSA of A7M08 is 1,736 Å^2^ with the heavy chain and light chain contributing 794 Å^2^ and 942 Å^2^, respectively, and have a nearly identical interacting profile as D24.1M01 (Fig S6). The affinity matured CDRL2, _49_FVY_51_ in D24.1M01 and _49_FTY_51_ in A7M08, sit above a small aliphatic surface formed by the EF- and FG-loops of P1 and the HI-loop of P2. The hydroxyl group of the germline Y49 could clash with the aliphatic interface between the EF- and HI-loops. The polar N51 in the germline V_H_2-70 could also clash with the aliphatic groove. The structure also showed a potential mechanism for the A7M08 Fab susceptibility to BC-loop mutations. V_H_2-70 encodes for a cysteine residue in the CDRH1. D24.1M01 contains a cysteine in the CDRH3 that forms a disulfide bond with the CDRH1 (Fig S7A) that could help to lock the CDRs into this orientation. A7M08 instead has a bulky tyrosine at the equivalent position (Y97) in the CDRH3 that would potentially make it more sensitive to mutations in the BC-loop. In conclusion, these structures show that V_H_2-70/V_L_λ1-40 mAbs bind to the HPV16 L1 with nearly identical epitopes and gives structural insight into the necessary affinity maturation of the CDRL2.

### V_H_4-34 with D3-16*02 bind FG-loops

The other frequently used IGHV was V_H_4-34, found in eight clonotypes (10 mAbs). Alignment of heavy chain sequences employing V_H_4-34 revealed a homologous sequence: Trp-Gly-Ser-Tyr-Arg (_100_WGSYR_100D_, in B25M05) in the CDRH3, which is predicted to be encoded by diversity region D3-16*02, and was present in five of eight (62.5%) clonotypes using V_H_4-34 (Fig 5A) compared to just four of 46 (8.7%) clonotypes using other V_H_ (p = 0.002). Antibodies using V_H_4-34 required the FGb loop for neutralization (Fig 5B). There were five mAb that utilized D3-16*02 that did not utilize V_H_4-34 that also required FGb either in the primary or secondary screens (not shown). Five of these 15 antibodies required the DE loop in addition to the FG loop for neutralization. Replacing the WGSYR sequence with five alanines in D25M07 resulted in a loss of neutralization activity (Fig 5C). When individual alanine substitutions were made, Y-to-A_100C_ showed the greatest loss of neutralization potency with an IC_50_ increase (from 1.6 +/- 0.2 pM for the mature antibody to 50.8 +/- 12.4 pM for the single amino acid substitution). Substitutions at the other four positions increased the IC_50_ modestly in comparison (W_100_), 2.6 +/1 0.6; G100, 3.9 +/- 0.9; S_100B_, 2.2 +/- 0.4; R_100D_, 6.2 +/- 1.3 pM). Thus, the V_H_4-34 heavy chain was frequently used in mAbs that recognize an epitope in the FGb loop for neutralization and tended to use D3-16*02 resulting in a conserved sequence in the CDR3 region (WGSYR) that was important for neutralizing activity.

**Fig 5.**
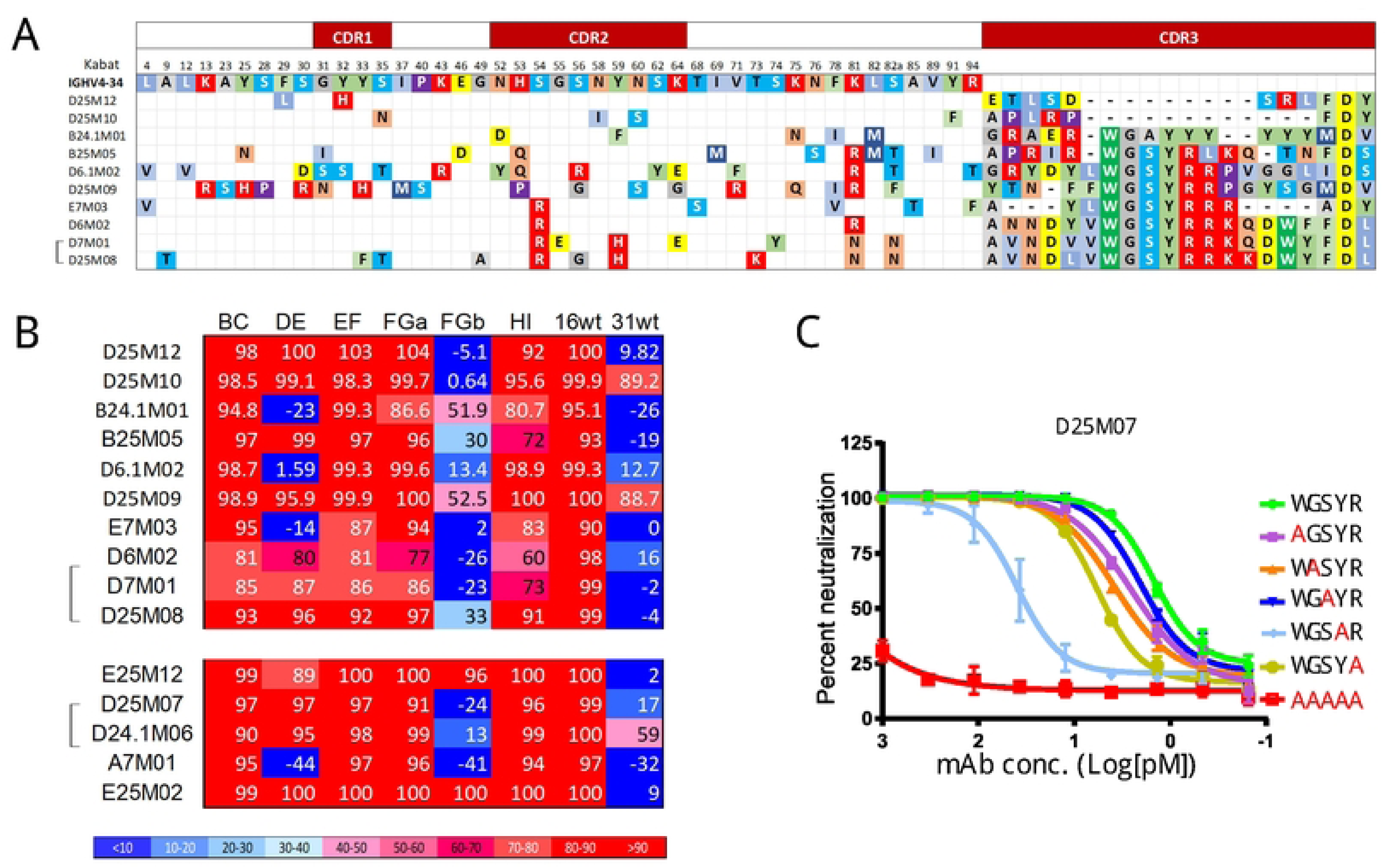
Comparison of mAbs utilizing IGVH4-34. A) Sequences alignment of mAbs using V_H_4-34 compared with the presumed germline sequence IGHV4-34*01. Dashes indicate a gap and empty cells indicate a residue the same as the germline. B) Neutralization results (percentages) for 10 mAbs in (A) and five mAbs that use IGHV4-34*01 but have other IGHV (below). C) Residues WGSYR in the CDRH3 region of D25M07 were substituted with alanines individually or with five alanine amino acids. Purified antibodies were serially diluted and tested for neutralization of psV16 along with unmutated D25M07.

### Structure of V_H_4-34 mAb in complex with L1 by cryo-EM

To investigate the binding mechanism of the V_H_4-34 mAbs, we chose two antibodies that had WGSYR in the CDRL3 region but had different patterns of neutralization (B25M05 and E7M03) however we could not obtain high resolution data for E7M03 (not shown). Cryo-EM was used to determine the structure of B25M05 Fab bound to the HPV16 L1 pentamer to a resolution of 2.9 Å (Fig 6A, Table S4, Fig S4). In contrast to the V_H_2-70/V_L_λ1-40 mAbs, only one B25M05 Fab bound to the pentamer (Fig 6B). The Fab bound to the top, interior of the pentamer at the interface between the DE-loops of P1 and P2 and the FG-loop of P1, with a total BSA of 941 Å^2^. On P1 the DE- and FG-loop are buried by 295 and 234 Å^2^, respectively, and the DE-loop of P2 is buried by 412 Å^2^ (Fig S6). B25M05 binds with a total BSA of 839 Å^2^, with 679 Å^2^ from the HC and 150 Å^2^ from the LC (Fig S5). The binding is mainly from the CDRH3, accounting for 72% of the total BSA, with some CDRH1 (8% of the total BSA) and CDRL1 (15% of the total BSA) (Fig 6C). The conserved WGSYR motif in the CDRH3 intercalates between the three variable loops (Fig 6D), displacing the DE-loop of P1, relative to the loop orientation in the non-bound protomers (Fig S7B). Y100C, which was shown to be most sensitive to alanine mutation, sits at the bottom of the pocket formed by the FG-loop of P1 and the DE-loop of P2. The FGb-loop of P1 makes substantial contact with _100B_SYR_100D_ residues of the CDRH3 and could explain why the V_H_4-34 mAbs are sensitive to mutations in the FGb-loops. Overall, this structure reveals a drastically different binding mechanism of the V_H_4-34 mAb to the HPV16 L1 pentamer and shows how the conserved WGSYR motif in the CDRH3 is crucial for binding.

**Fig 6.**
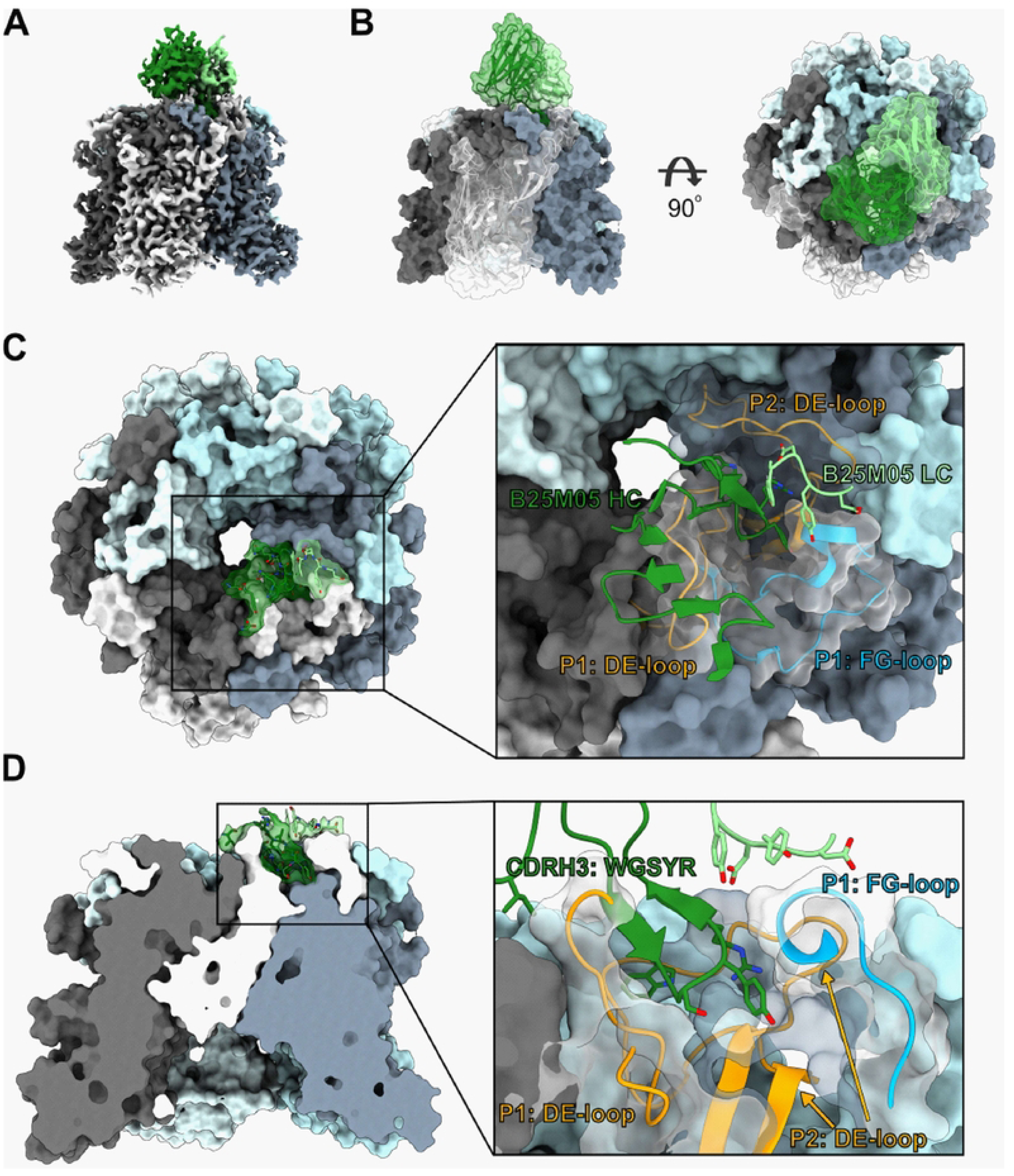
Structure of IGVH4-34 antibody B25M05 bound to HPV16 L1. A) Cryo-EM map of B25M05 Fab bound to the HPV16 L1 pentamer. L1 protomers are colored in shades black, white, and blue. The heavy and light chains of B25M05 are colored dark green and light green, respectively. B) A surface representation of the structure in two orientations. One protomer and the Fab is shown with a transparent surface with a cartoon representation. C) Top view of the binding interface of B25M05. The insert shows the three L1 loops involved in binding. D) Side view of the L1 pentamer cut out to show the binding pocket of the B25M05 CDRH3. Insert shows the WGSYR motif and how it binds in a pocket formed by the two protomers.

## Discussion

As expected, most HPV16 mAbs were type specific and required sequences on one or more surface loops for neutralization activity. All surface-exposed loops on HPV16L1 were required for neutralization by one or more mAbs suggesting that much of the surface of HPV16 can be targeted by antibodies. In agreement with previous studies, we found that the FG-, HI- and DE- loops were frequently required for neutralization by mAbs. More than half of the antibodies studied here required FG for neutralization and DE- and HI-loops were required for neutralization by more than 20% of the mAbs tested. We could not mutate all surface-exposed residues without disrupting infectivity, therefore the methods used here likely underestimate the antibody diversity. The broad antibody diversity observed here helps to account for the nearly universal seroconversion to prophylactic HPV vaccination and the high proportion of neutralizing mAbs may explain the high efficacy of these vaccines ^40^. Three mAbs required sequences on the C-terminus for neutralization, a result seen previously ^14,17^.

HPV16-HPV31 cross-reactive serum antibodies are commonly induced by the bivalent and quadrivalent HPV vaccines and both vaccines have shown protection against HPV31 and other cross-reactive non-vaccine types ^19,41^. Previous studies have demonstrated that sera from HPV vaccinees recognize cross-reactive neutralizing epitopes on the DE- and FG-loops ^15,16^ ^19,41^. Bissett et. al. ^16^ demonstrated that sera from HPV vaccinees recognize cross-reactive neutralizing epitopes on the DE- and FG-loops. The six HPV31 cross-reactive antibodies we identified all required the C-terminal portion of the FG-loop for neutralization (Fig 2). Interestingly, there were four mAb of one clonotype that had similar HPV16 neutralization titers but variable levels of HPV31 cross-reactivity (Fig 2). Swapping heavy and light chains among these mAbs revealed that the light chain conferred most of the psV31 cross-neutralization activity (Fig 2). Among the 54 clonotypes described here, the six cross-reactive antibodies had a higher number of putative amino acid changes in the heavy and light chains (12.70 +/- 7.61) compared to the non-cross-reactive antibodies (9.48 +/- 4.90; p = 0.048) and in the clonotypic family, the cross-reactive mAbs had more of somatic mutations in their heavy and light chain variable regions than the non-cross-reactive mAbs (not shown) implying that cross-reactive memory B cells were derived from non-cross-reactive precursors. This finding is reminiscent of the development of broadly neutralizing antibodies to influenza and HIV ^42,43^ that evolve from more type-specific precursor B cell ^42,43^.

The genetic characteristics of antibodies that target specific epitopes have been studied for other viruses ^31,33,34,44–46^. This is the first study to identify recurrent features of HPV specific mAbs that correlated with patterns of neutralization. We found that V_H_2-70 frequently paired with the V_L_λ1-40 and those antibodies had characteristic mutations in the CDRL2 region (Fig 3C). Pairing of unique heavy and light chains has been observed among recurring antibodies following viral infection and vaccination ^31,47–49^. Recurrent Ebola virus antibodies used V_L_λ1-40 in combination with IGHV3–15. ^47^ ^31,48,49^. Those antibodies also had a characteristic pattern of mutations in which the CDRL2 serine was substituted with asparagine or threonine, a pattern not observed here. Here we found the CDRL2 of antibodies that paired V_H_2-70/ V_L_λ1-40 was important for HPV16 neutralization. Substitutions with the predicted germline sequence in A7M08 resulted in an antibody with greatly reduced neutralization capacity. Even a single amino acid change from the germline CDRL2 sequence to the residue found in the mature mAb, improved neutralization activity more than two logs and even a conservative substitution (YGN to FGN) improved neutralization over one log compared to germline. Similar results were observed for four of the other six V_H_2-70/V_L_λ1-40 mAbs as swapping the mature CDRL2 with the germline CDRL2, increased their IC_50_s, albeit to a lesser extent than for A7M08 (Table S3). Although, the psV16 neutralization activity of two mAbs (B25M02, D25M03) were not similarly affected by substitutions in the CDRL2, we showed that in the context of the germline light chain, amino acid substitutions in the CDRL2 significantly improved neutralization activity (Fig S3). These results support the importance of the V_L_λ1- 40 CDRL2 for HPV16 recognition and neutralization, as implied by the apparent convergent evolution for the sequences in this region however, cryo-EM revealed that V_L_λ1-40 contacted L1 with residues in CDRL1, CDRL3 and framework regions in addition to CDRL2 residues (S5 Fig).

The structures of the two antibodies with paired 2-70/V_L_λ1-40, bound to capsomers revealed nearly identical footprints (Fig 4). It was difficult to discern which residues were required for neutralization for antibodies in this set using cpsV with, all but A7M08, able to neutralize the entire panel of cpsV in the first neutralization screen (Fig 3D). This likely resulted from these antibodies having similar overlapping epitopes dispersed across the five loops and our assay was not sensitive enough to detect minor disruptions in neutralization potency when only one loop had amino acid substitutions. The footprint of D24.1M01 and A7M08 overlaps the published footprints for other HPV16 monoclonal antibodies including H16.V5 ^21–23^. D24.1M01 and H16.V5 both footprint onto identical residues on all five loops (P55, A139, N181, P182, V267, K278, N285, I348, T358)^22^. These antibodies have been described as binding to the top fringe of pentamers to multiple loops, primarily DE, FG and HI ^21^. The majority of papillomavirus binding antibodies described previously share this pattern of binding, thus this appears to be the immunodominant region of the virus and could be described as a public epitope, although, given the diversity of antibodies that bind here, this region might expose several overlapping epitopes.

The other set of antibodies focused on here were antibodies using IGHV4-34. There were ten mAbs using IGHV4-34 representing eight clonotypes that paired with seven different IGLV but surprisingly, five paired with the same diversity region gene segment (D3-16*2, same reading frame). Because diversity gene segments encode part of the CDRH3 region of it was unsurprising to find a conserved sequence (WGSYR) in this region. Using alanine scanning we showed that this conserved sequence, in particular the tyrosine was important for neutralization activity (Fig 5). Diversity gene segments ^50^ have been found to be important for broadly neutralizing influenza antibodies and hydrophobic antibody residues were found to be important for those interactions ^50^. Although all mAbs with WGSYR in the CDRH3 required the C-terminal half of the FG-loop for efficient neutralization, they did not recognize an identical epitope, with four requiring the DE-loop for neutralization, and two having significant cross-reactivity with HPV31. The cyro-EM structure of B25M05 was strikingly different from A7M08 and D24.1M01. Instead of five Fabs per capsomer, B25M05 bound one Fab per capsomer at the pentamer apex. This structure has been observed with other papillomavirus binding antibodies suggesting that this is also an immunodominant public epitope.

One limitation of this study is that we could not make mutations to all residues on the surface loops. Using cpsV to characterize neutralization epitopes relies on particles folding into native conformations (an approach also taken by Bissett et. al. ^16^) but limits us to cpsV that retain infectivity. Thus, the BC-loop has only two substitutions and one deletion across the 11-residue loop compared with 10 substitutions across the 14 residue FGb-loop. This certainly reduced the sensitivity for detection of BC specific mAbs compared with FGb specific mAbs. Additionally, the C-terminal portion of DE- and HI-loops could not be mutated without impairing infectivity. The mouse mAb H16.V5 was tested in this assay and FG-loop was confirmed to be required for neutralization, but this method did not detect the HI-loop that is a known binding site of this antibody (not shown) ^12,24,35^. Another limitation is the arbitrary nature of the thresholds chosen to call an antibody dependent on a certain loop. Thresholds were set to cluster antibodies with similar neutralization profiles together. We could then assess if antibodies that shared neutralization profiles also shared molecular features.

In summary, we have shown that all L1 surface loops are targeted by human antibodies with the FG loop containing a dominant neutralizing epitope. Although this set of antibodies was modest in size, we found two types of antibodies that appeared to be over-represented and bound to the virus in strikingly different fashions. Importantly, both types of antibodies were found in each of the four participants in this study, suggesting that these may be commonly induced antibodies that recognize public epitopes.

## Materials and methods

### Cell culture

Cells (293TT generously provided by John Schiller, National Cancer Institute, USA) were grown in DMEM (Thermo Fisher Scientific, Waltham MA) with 10% Bovine serum albumin (Corning, Corning, NY)) and supplements as previously described ^51^. Pseudovirus were produced and purified as described by Buck et al ^51^ except that four T225 cm^2^ plates (Corning) were transfected, each with 57 μg DNAs (cpsV and pYSEAP plasmids) with 171μL TransIT®-293 reagent (Mirus Bio LLC, Madison WI). All psV were purified on Optiprep gradients (Millipore Sigma Burlington, MA). Monoclonal antibody cloning and expression were previously described ^52^Cells (293TT generously provided by John Schiller, National Cancer Institute, USA) were grown in DMEM (Thermo Fisher Scientific, Waltham MA) with 10% Bovine serum albumin (Corning, Corning, NY)) and supplements as previously described ^51^. Pseudovirus were produced and purified as described by Buck et al ^51^ except that four T225 cm^2^ plates (Corning) were transfected, each with 57 μg DNAs (cpsV and pYSEAP plasmids) with 171μL TransIT®-293 reagent (Mirus Bio LLC, Madison WI). All psV were purified on Optiprep gradients (Millipore Sigma Burlington, MA). Monoclonal antibody cloning and expression were previously described ^52^.

### Mutagenesis of psV and mAb

Sequences of HPV16 L1 targeted for substitution were identified by alignment of HPV16, HPV18 and HPV2 L1 protein sequences. Residues constituting loops were as defined by Chen et al ^10^. All mutagenesis was performed on the p16sheLL plasmid (Addgene, Cambridge, MA) using oligonucleotide primers (Integrated DNA Technology Inc (IDT), San Jose, CA) and In Fusion® cloning technology (Clontech Inc., Mountain View, CA). Sequence changes were verified by Sanger sequencing (Genewiz, South Plainfield, NJ). Initially all loops were substituted with HPV18 sequence but not all constructs produced infectious cpsV. Non-infectious cpsV were remade using HPV2 sequences and if those cpsV were non-infectious, the process was repeated substituting only a portion of each loop with HPV18 or HPV2 sequences until infectious cpsV were obtained with mutations on each of the five surface loops. Mutation of antibodies in the AbVec plasmids ^53^ were made using similar methods. Mutations to CDRL2 of V_L_λ1-40 in multiple antibodies and the CDRH3 of V_H_3-34 of D25M07 were created using the methods described above or using the Q5® Site-Directed Mutagenesis Kit (New England, BioLabs, Ipswich, MA) with primers and sequencing performed as above. Designation of CDR regions and numbering was adopted from IGMT ^54^. The entire V_L_λ1-40 sequence (accession M94116 with modifications to the CDRL3 region to match B25M02) was codon modified and synthesized (IDT) and cloned into Abvec-hIglambda (FJ517647.1) kindly provided by Patrick Wilson’s lab (University of Chicago).

### Neutralization assays

To determine the optimal concentration of each cpsV, 3X10^5^ 293TT cells were plated in the wells of a 96 well flat bottom tissue culture plate (Corning) in 100 μl of medium. Approximately four hours later cpsV were serially diluted across a 96 well polypropylene plate (USA scientific, Orlando, FL) excluding the outer wells. After incubation at room temperature for one hour, 100 μl of diluted cpsV was used to infect the 293TT cells (adding media only to the last row) and transferred to a 37°C incubator (5% CO_2_) for three days. Supernatants (30 μl per well) were carefully removed and transferred to a 96 well ELISA plate (Fisher Scientific International, Hampton, NH). Secreted alkaline phosphatase was detected using phosphatase 104 substrate (Millipore Sigma) dissolved in 0.1M Carbonate buffer (pH9.6) with 10 mM MgCl_2_ (100 μl per well). After a 30-minute incubation in the dark, plates were read on a Synergy H4 plate reader (BioTek, Winooski, VT) at 405nm. Dilutions of cpsV that resulted in an absorbance of approximately 1.5 were determined to be the optimal for maximizing signal/noise ratios.

Primary mapping experiments were performed using a concentration of mAb 100-fold higher than the IC_50_ determined using psV16. Pseudovirus were diluted to the optimal concentration and mixed with diluted mAb in triplicate. The remainder of the assay was conducted as above. Background alkaline phosphatase was determined for each plate and was the average of six uninfected wells. Percent neutralization was calculated for each cpsV [1-((cpsV + mAb) – background)/ ((cpsV)-background)]. Repeat testing of 10 antibodies was highly reproducible (median coefficient of variation = 2.9%).

Human mAbs that neutralized all cpsV more that 75% were retested in a secondary screen using one row of a 96 well plate for each mAb/psV combination. Antibodies were serially diluted 1:3 across a plate starting at a dilution 100-fold higher than the IC_50_ for psV16, mixed with cpsV with subsequent steps conducted as above. To compute IC_50_ values the percent neutralization values for each well were fit to a four-parameter logistic regression equation (4-PL) using GraphPad Prism7.0 (GraphPad Software Inc, San Diego, CA).

Neutralization assays using mutated mAbs began at 100 nM with 3-fold serial dilutions. Antibodies that did not titrate down to background were retested starting at a lower concentration (10 nM). Antibodies that failed to neutralize at 100 nM (>50% neutralization) were classified as non-neutralizing. All antibodies were tested a minimum of three times and the median coefficient of variation for repeated testing was 20.5% for the IC50 values. Most experiments used purified mAb with known concentrations but for the neutralization experiments testing the c-terminal mutated psV some of the antibodies were cell culture supernatants. For those experiments antibody dilutions were used to calculate the EC50 rather than antibody concentrations.

### Statistical analysis

Clonally related antibodies (clonotypes) were defined here as being derived from the same subject, having the same heavy and light chain variable regions (as assigned by IMGT-V Quest), with CDRH3 regions having the same length and greater than 70% amino acid sequence homology. All statistical analyses and heat maps were conducted on GraphPad Prism. Fishers’ exact tests were used to determine if mAb molecular features were associated with patterns of neutralization or other molecular features. Mathematic calculations were conducted on EXCEL 365 (Microsoft corporation, Redmond WA, USA). Kabat numbering: http://www.abysis.org/abysis/

### Cloning and deletion construct for HPV16-L1

The full-length HPV16-L1 clone in pGEX-2T was used as parent clones for generating the deletion mutant. The L1 protein was expressed fused to GST protein tag in pET28b vector. We used a pair of forward and reverse primers to amplify the clone using the standard PCR method, except for the deleted regions. The regions deleted were N-terminal 9 residues, C-terminal 31 residues, and helix 4 (residues 404– 436), known to be required for HPV16 L1 virus particle assembly (15). The PCR fragment with NdeI and BamHI restriction sites incorporated into both primers was digested with the same enzyme for ligation. DNA sequencing was used to confirm the deletion mutant.

### HPV16L1 protein purification

The expression and purification of HPV16L1 protein was done as described previously #2499} Briefly, the E. coli (pLysS) cell culture was grown to the O.D. of 0.3 at 37 C, induced with 0.5 mM IPTG, and the protein was allowed to overexpress overnight at room temperature. The cells were pelleted and resuspended in lysis buffer having 50 mM Tris-HCl, pH 8.0, 0.2 M NaCl, 1 mM dithiothreitol (DTT), 1 mM EDTA, and 10 mM phenylmethylsulfonyl fluoride. The cells were lysed by a sonication cycle of 30sec on and 30sec off for 30 min.

Further, 5mM MgCl2 and 1mM ATP was added to the lysate and incubated for 1hr at room temperature on rocker. The lysate was centrifuged for 1hr at 10000 rpm and the supernatant was passed through GST bead column by gravity flow. The GST beads were washed with lysis buffer to remove the contaminating proteins and the L1 protein was eluted with lysis buffer having 10mM glutathione. The eluted L1 protein was treated overnight at 4 °C with thrombin enzyme (Sigma; T6634) to remove the GST tag. The cleaved L1 protein further passed through GST beads to remove the GST protein and to get a pure L1 protein. The eluted L1 protein was purified by HiLoad 16/600 Superdex 200 pg column.

### Expression and purification of Fabs

The plasmid encoding Fabs were expressed in 500ml of HEK293-EBNA cells were cultured to a density of 1 million cells/mL and transfected with 125 μg each of Fabs Heavy and light chains using 1 mg of PEI. Cultural supernatant was harvested 6 days after transfection. Fabs were purified using Ni-NTA resin (Takara) and eluted using 5mM HEPES, 150 mM NaCl, and 250mM imidazole. The fabs were further purified using HiLoad 16/600 Superdex 200 pg column.

### Cryo-EM of complexes

The HPV16L1-D24M01Fab complex was prepared by mixing HPV16L1 with Fab fragments in the ratio of 1:1.5 and then incubated at room temperature for 2 h. The incubated complex was further purified using HiLoad 16/600 Superdex 200 pg column. Aliquots of the purified complexes were loaded and vitrified on Ultrathin Carbon Film on Lacey Carbon Support Film (400 mesh, Cu), (Tedpella) in an FEI Mark IV Vitroblot. 2,332 movies were acquired using Titan Krios microscope (ThermoFisher) at a voltage of 300 keV equipped with Falcon 3EC direct electron detector (ThermoFisher) at counting mode at 96,000x (1.054 Å pixel size). Automated data collection was done using EPU (ThermoFisher) varied the defocus range between −0.8 to −2.2 μm. All movies were collected using an exposure time of approximately 5 s for a total dose of 50 e−/Å2 fractionated over 50 frames.

The HPV16L1-A7M08Fab complex was prepared the same way as the D24M01 complex. Sample was loaded at a concentration of 0.25 mg/mL with 0.04% Dodecyl-β-D-maltoside (DDM) onto UltrAuFoil R2/2 on 200 mesh (Ted Pella) using a Vitrobot Mark IV (ThermoFIsher). Grids were blotted for 6 s with a blot force of 2 at 4 °C and 100% humidity. Data was collected on a 200 kV Glacios microscope (ThermoFisher) with a K3 direct electron detector (Gatan) at a magnification of 36,000x (1.122 Å per pixel). Initial analysis showed orientation bias at 0° so a total of 2,372 movies were collected, 560 at 0°, 1,237 at 30°, and 575 at 45° tilt using SerialEM ^55^ for automated collection with a defocus range of -1.5 to -2.2 μm with exposures over 6 s.

The HPV16L1-B25M05Fab complex was prepared in the same way as the D24M01 complex. Sample was loaded at a concentration of 0.6 mg/mL with 0.05% DDM onto UltrAuFoil R2/2 on 200 mesh (Ted Pella) using a Vitrobot Mark IV (ThermoFIsher). Grids were blotted for 4 s with a blot force of 2 at 4 °C and 100% humidity. A total of 1,079 movies were collected without tilt on a 200 kV Glacios microscope (ThermoFisher) with a K3 direct electron detector (Gatan) at a magnification of 36,000x (1.122 Å per pixel) using SerialEM ^55^ for automated collection with a defocus range of -1.8 to -2.2 μm with 100 exposures over 6 s.

### Cryo-EM Data processing

For the D24M01 data, alignment and CTF estimation of aligned micrographs were carried out with Motioncor2 ^56^. Processed micrographs were imported into cryoSPARC (v3.1) ^57^. Particles were selected using blob picker and extracted with a box size of 220 px. Following two rounds of 2D classification (150 classes and 100 classes), there were 258,311 particles. Two *ab-initio* reconstructions were generated and following heterogeneous refinement with C5 symmetry, one showed much higher resolution (3.55 Å and 5.35 Å). The higher resolution reconstruction was further processed using Local CTF refinement and non-uniform refinement with 188,743 particles. The final 3D density map resolution was estimated based on the gold-standard FSC (GSFSC) curve with a cut-off of 0.143 (GSFSC_0.143_) to 3.0 Å. Information for the 3D reconstruction can be found in Table S4 and Figure S4.

For the A7M08 data, motion correction was performed using WARP and further processed using cryoSPARC (v4.4) ^57^. Micrographs were curated down to a total of 1,584 micrographs, 449 at 0°, 937 at 30°, and 198 at 45°. Template picker was used to pick particles using the map of the D24M01 with HPV16 L1 to generate templates and particles were extracted using a box size of 320 px. Following three rounds of 2D classification (150 classes, 100 classes, and 50 classes, there were 989,215 particles. A single *ab-initio* reconstruction was generated and 3D classing was used to further select particles. From this, 301,713 particles were used to generate the final reconstruction to a GSFSC_0.143_ resolution of 2.69 Å. Information for the 3D reconstruction can be found in Table S4 and Figure S4.

For the B25M05 data, motion correction was performed using WARP and processed using crySPARC (V4.4) ^57^. Micrographs were curated down to 753 and particles were picked using blob pickers and extracted with a box size of 320 px. A single *ab-initio* reconstruction was generated but refinement failed to give a high-resolution map. Particles from the best homogeneous refinement were symmetry expanded with C5 symmetry and 3D classified into 5 classes. The best refined map was moved forward and duplicate particles were removed. The final reconstruction was generated using 83,711 particles with C1 symmetry to a final reconstruction GSFSC_0.143_ resolution of 2.90 Å. Information for the 3D reconstruction can be found in Table S4 and Figure S4.

### Model building and refinement

The template model was fitted into sub-particle reconstruction density maps using ChimeraX ^58^. The models were then corrected and adjusted manually by Isolde ^59^ in ChimeraX and COOT ^60^ and automatically in real_space_refinement in Phenix ^61^. The model statistics is summarized in Table S4. PISA server (www.ebi.ac.uk/pdbe/pisa) was used to analyze the L1-Fab interaction.

## Acknowledgements

We thank Duncan Ralf and Eric Matsen for helpful discussions. This research was supported by the Electron Microscopy Shared Resource, RRID:SCR_022611, of the Fred Hutch/University of Washington/Seattle Children’s Cancer Consortium (P30 CA015704). A portion of this research was supported by NIH grant R24GM154185 and performed at the Pacific Northwest Center for Cryo-EM (PNCC) with assistance from Harry Scott.

## Author Contributions

### Conceptualization

Erin Scherer, Joseph Carter, Denise Galloway, Marie Pancera

### Data curation

Erin Scherer, Joseph Carter, Robin Smith

### Formal analysis

Joseph Carter, Nick Hurlburt

### Funding acquisition

Marie Pancera, Denise Galloway

### Investigation

Joseph Carter, Nick Hurlburt, Erin M. Scherer, Suruchi Singh, Justas Rodarte, Robin A. Smith, Peter Lewis, Rachel Kinzelman, Jacqueline Kieltyka, Madelyn Cabãn, Gregory Wipf

### Methodology

Erin Scherer, Joseph Carter, Marie Pancera

### Project administration

Denise Galloway

### Resources

Marie Pancera, Denise Galloway

### Supervision

Marie Pancera, Denise Galloway

**Validation:**

### Writing – original draft

Joseph Carter, Nick Hurlburt

### Writing – review & editing

Joseph Carter, Nick Hurlburt, Marie Pancera, Denise Galloway

## Supporting information

**Table S1. Cross reference for antibodies with previous nomenclature.** Antibodies in this manuscript (columns 1,3) are cross referenced to identifiers used in Scherer et. al. 2018 [22]. The first letter in the current ID represents the study subject, the following number is the month at which the sample was collected. If the number is a decimal it was collected one week after the vaccine dose. The letter M or P indicates the source was a memory B cells or plasmablast and last two digits were added to create unique identifier.

**Table S2. Screen for determining which loops were required for neutralization**. These are the same data shown in Fig. 1B however in this figure the percent neutralization is shown. Antibodies were tested in triplicate in neutralization assays against sixcpsV, psV16 and psv31 using a concentration of antibody of 100 fold higher than their IC50 versus psV16. Each cpsV had amino acid substitutions on one of the five surface loops of HPV16 as indicated in Fig. 1A. The value is percent neutralization: setting psV with no antibody equal to zero and no psV equal to 100%. Clustering was performed manually to clonotypes together. Clonotypes are indicated by brackets on left.

**Table S3. Measuring the importance of the CDRL2 region for neutralization of psV16.** The germline CDRL2 sequence was cloned into the light chains for each mAb listed. The modified light chains were coexpressed with their cognate heavy chain and used in neutralization assays with psV16. The fold differences are compared with results using the mature light chain (see sequence Fig 3C) for each.

**Table S4. Cryo-EM data collection and refinement statistics.** Data collection information for the cryo-EM data and statistics from refinement of the models. MolProbity ^62^ and EMRinger ^63^ were used for analysis of the quality of the final model.

**Figure S1. Most HPV16 binding antibodies neutralize.** 148 hmAb, including the 70 mAb included in this study were tested for HPV16 neutralization and binding. Antibodies that failed to neutralize were given an arbitrary IC50 of 100 nM. There were four mAb (circled) that had significant binding but failed to neutralize. The p value is from linear regression, indicating that the slope of the line is statistically significantly not equal to zero.

**Figure S2. Human mAbs potently neutralize HPV16 in vitro.** A histogram showing the distribution of HPV16 IC50 neutralization values for the 70 hmAbs used in this study.

**Figure S3. Importance of CDRL2 for B25M02 neutralization.** Because changing just the CDRL2 region in the otherwise mature B25M02 didn’t significantly reduce the neutralization we wanted to test the neutralization potency of the germline B25M02 light chain with just the CDRL2 mutated – see sequences – panel A. B25M02 heavy chain was coexpressed with either the fully mature B25M02 light chain, fully germline IGLVl1-40 light chain or the germline light chain with the mature (HVF) CDRL2. Each data point (B) is an IC50 vs psV16, determined from a titration curve. *** = p<0.001. ns = not significant.GL = germline seq.

**Figure S4. Data processing of cryo-EM data.** Cryo-EM reconstructions of D24.1M01 Fab bound to HPV16L1 (left), A7M08 Fab bound to HPV16L1 (middle), and B25M05 Fab bound to HPV16L1 (right). Representative micrographs, representative 2D class averages, final resolved map colored based on local resolution estimation, and FSC plot with resolution estimation based on GSFSC_0.143_ are shown.

**Figure S5. BSA of mAbs D24.1M01, A7M08, and B25M05 binding to HPV16 L1.** Buried surface area (BSA) of each mAb amino acid upon binding with HPV16 L1. Values were determined using the PDBePISA server ^64^. Interactions with protomer 1 are outlined in gray and interactions with protomer 2 are outlined in black.

**Figure S6. BSA of HPV16 L1.** Buried surface area (BSA) of the amino acids of HPV L1 upon binding by D24.1M01 or B25M05. Values were determined using the PDBePISA server ^64^. Heavy and light chain interactions are differentiated in dark and light colors. Amino acids from protomer 1 are solid colors and amino acids from protomer 2 have a checkered pattern.

**Figure S7. Importance of disulfide bond linking D24.1M01 CDRH1 and CDRH3 and movement of DE-loop in B25M05 binding.** A) Cartoon diagram of the overlayed D24.1M01 and A7M08 structures. D24.1M01 has a disulfide bond linking the CDRH1 and CDRH3 but in A7M08 the CDRH3 contains a tyrosine. B) Cartoon diagram showing the movement of the DE-loop between unbound and B25M05 bound L1 protomers. Only the DE-loop undergoes rearrangement the FG-loop is unchanged upon binding.

